# Mechanical mapping of spinal cord development and repair in living zebrafish larvae using Brillouin microscopy

**DOI:** 10.1101/181560

**Authors:** Raimund Schlüßler, Stephanie Möllmert, Shada Abuhattum, Gheorghe Cojoc, Paul Müller, Kyoohyun Kim, Conrad Möckel, Conrad Zimmermann, Jürgen Czarske, Jochen Guck

**Author notes:** these authors contributed equally to this work.

## Abstract

The mechanical properties of biological tissues are increasingly recognized as important factors in developmental and pathological processes. Most existing mechanical measurement techniques either necessitate destruction of the tissue for access or provide insufficient spatial resolution. Here, we show for the first time a systematic application of confocal Brillouin microscopy to quantitatively map the mechanical properties of spinal cord tissues during biologically relevant processes in a contact-free and non-destructive manner. Living zebrafish larvae were mechanically imaged in all anatomical planes, during development and after spinal cord injury. These experiments revealed that Brillouin microscopy is capable of detecting the mechanical properties of distinct anatomical structures without interfering with the animal’s natural development. The Brillouin shift within the spinal cord increased during development and transiently decreased during the repair processes following spinal cord transection. By taking into account the refractive index distribution, we explicitly determined the apparent longitudinal modulus and viscosity of different larval zebrafish tissues. Importantly, mechanical properties differed between tissues *in situ* and in excised slices. The presented work constitutes the first step towards an *in vivo* assessment of spinal cord tissue mechanics during regeneration, provides a methodical basis to identify key determinants of mechanical tissue properties and allows to test their relative importance in combination with biochemical and genetic factors during developmental and regenerative processes.

## Introduction

In response to a mechanical stimulus, many cell types, including neurons and glia, change their properties and behavior^1–5^. This includes morphological changes^6–8^ as well as alterations of migration^9^, proliferation^10^ and differentiation^7^. The cellular susceptibility to mechanical signals is termed mechanosensitivity and has been the subject of recent studies documenting the involvement of mechanical signals in developmental and pathological processes. Axonal growth during optic tract development in *Xenopus laevis*, for instance, is guided by temporally changing stiffness gradients in adjacent brain tissue^11^. Microglial cells and astrocytes in rat brains display an activated phenotype and increase expression of inflammatory genes and proteins, which is indicative of reactive gliosis, when exposed to a mechanical stimulus that deviates from their physiological mechanical environment^12^. Oligodendrocyte precursor cells increase their expression of myelin basic protein and display an elaborated myelin membrane on stiffer substrates as compared to more compliant substrates indicating a preferred mechanical environment for differentiation^7^. Among others, these examples demonstrate an intricate interplay between the mechanosensitivity of distinct neural cell types and the mechanical properties of their respective microenvironments. *In vivo*, these microenvironments are formed by the surrounding nervous tissue whose mechanical properties are thought to be determined by the combined material properties of neighboring cells, extra cellular matrix and the degree of cross-linking^13^. As these determinants of tissue mechanics change during development and after pathological events, concomitant changes of mechanical tissue properties and their direct involvement in a wide range of central nervous system conditions and diseases are likely^14–17^.

Severe injury to the mammalian spinal cord prompts a biochemical signaling cascade that eventually leads to the formation of a glial-fibrotic scar. The scar tissue has not only been shown to prevent axonal regrowth across the lesion site due to its biochemical composition^18, 19^, but has also been proposed to act as a mechanical obstacle^19, 20^. Unlike mammalian systems, in which the loss of motor and sensory function associated with traumatic spinal cord injury is permanent, zebrafish are able to restore motor function and repair spinal cord tissue even after complete spinal cord transection^21^. In adult zebrafish, functional recovery correlates with an orchestrated cellular response that includes proliferation^22–25^, migration^23, 25, 26^, differentiation^22, 25, 27^and morphological changes^22, 23^. Radial glial cells, for example, proliferate and give rise to new motor neurons that mature and integrate into the spinal circuitry^22^. Descending axonal projections severed by the injury initiate regrowth and eventually traverse the injury site^25, 28^. These regenerative events take place in a time interval of approximately six to eight weeks post-injury^22, 25^. Larval zebrafish have been shown to share the regenerative capacity with their adult counterparts, as they too respond to spinal cord transection by proliferation, neurogenesis, axonal regrowth and functional recovery^29–32^, yet with a much shorter recovery time^29, 31, 33^.

In light of the aforementioned examples of neural mechanosensitivity and stiffness changes of nervous tissue during development and disease, we submit that the mechanical characterization of zebrafish spinal cord tissue poses an intriguing addition to biochemical analysis and might help to elucidate the role of mechanical signaling during spinal cord development and successful spinal cord repair.

To quantify the mechanical properties of biological samples, a variety of standard measurement techniques has been developed^34, 35^. Micromanipulation of nanoparticles and cells^36^, ferrofluid microdroplets^37^, optical tweezers^38^, traps^39^ and stretchers^40^, micropipette aspiration^41^ and cell shape analysis^42^ allow the local measurement of mechanical properties of single cells or tissues at a molecular to cellular scale. Parallel plate measurements and tissue shape analysis^43^ provide mechanical data for bulk tissues. However, only a fraction of the techniques suitable for *in vivo* measurements of mechanical tissue properties allows the spatial mapping of the intrinsic mechanical properties of whole native tissues^44^. These include e.g. atomic force microscopy (AFM)-based indentation^45–47^, sonoelastography and magnetic resonance elastography^48^. While the former necessitates direct physical contact between indenter and sample, and can therefore only measure the surface of a dissected, possibly sectioned tissue, the latter lack spatial resolution in the range relevant for cellular mechanosensitivity and required for samples with small dimensions such as zebrafish larvae.

Brillouin spectroscopy allows measurements of mechanical properties and is widely used in the material sciences. The technique is based on the inelastic Brillouin scattering process between incident photons and mass density fluctuations (acoustic phonons) due to inherent, travelling sound waves in the sample (see. Methods)^49, 50^. It gives access to the phonon’s properties which are related to the viscoelasticity of the sample, e.g. the longitudinal storage and loss moduli. With the emergence of virtually imaged phased array (VIPA) based spectrometers in the last decade, this technique has become applicable for the point-wise measurement within living biological samples to build up a three-dimensional image with diffraction-limited resolution^51^. Brillouin microscopy has so far been applied to human cornea^52–54^, murine carotid arteries^55^, ruminant retina^56^, rabbit bone tissue^57^ and zebrafish embryos^58–60^. Here, we exploit the optical transparency and high regenerative capacity of zebrafish larvae to systematically explore the *in vivo* mechanical properties of larval tissue during development and repair using a custom-built Brillouin microscopy setup.

Our Brillouin microscope (Fig. 1) is based on a two-stage VIPA interferometer design from Scarcelli et. al.^51, 61–63^. As illumination source we use a frequency modulated diode laser (Toptica DLC TA PRO 780) emitting at 780.24 nm whose central frequency is stabilized to the D2 transition of rubidium ^85^Rb. For this purpose, a probe beam from the laser is transmitted through a Rb molecular absorption cell. The intensity behind the cell is measured by a reference photodetector and used as input signal for the lock-in frequency stabilization. The laser spectrum is cleaned using a Bragg grating (Ondax NoiseBlock) to increase the laser mode to amplified spontaneous emission (ASE) ratio by 10 dB to over 65 dB. While this ASE ratio already allows *in vivo* measurements in zebrafish larvae (shown in Figs. 2 to 5), some Brillouin spectra from positions with strong reflections inside the specimen cannot be evaluated (See Supplementary Note 1). This can be mitigated by a further suppression of the ASE. We therefore implemented a Fabry-Perot interferometer (FPI) in the illumination path whose cavity length is stabilized to the central laser frequency. For this purpose, the light reflected at the FPI is guided to a photodetector. The acquired intensity signal is then used as input signal for an adjusted Pound-Drever-Hall control scheme^64^. The FPI leads to a reduction of the laser mode to ASE ratio by 20 dB to approximately 85 dB. This improvement yields clean spectra for measurements that were previously affected by strong reflections. The improved setup was used for the measurements shown in the last two figures and all Brillouin measurements shown in the Supplementary Material.

**Figure 1.**
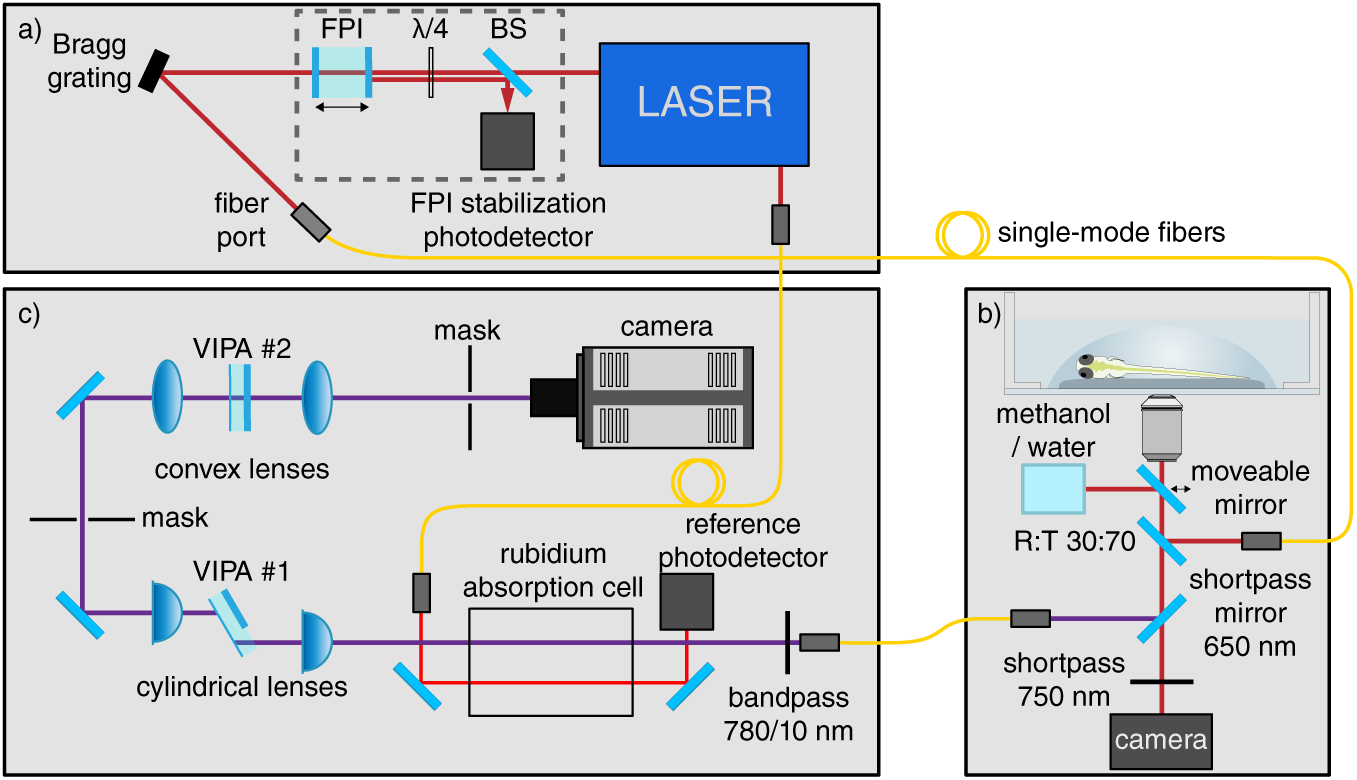
Scheme of the custom-built confocal Brillouin microscope. The setup consists of a) a diode laser to illuminate the specimen, b) a confocal microscope and c) the Brillouin spectrometer. The light inelastically scattered in the specimen is collected in backscattering direction and coupled into a two-stage VIPA spectrometer including a molecular absorption cell filled with Rubidium. The spectrogram is acquired with an sCMOS-camera and evaluated using a self-written MATLAB program. BS = polarizing beam splitter.

The laser light is then coupled into a single-mode fiber and guided to a custom-built inverse confocal microscope. The confocal microscope is based on a Zeiss Axiovert 200M microscope stand equipped with a translation stage. The light emitted from the illumination fiber is collimated, guided to the objective and focused into the specimen with a total power of 10mW. Alternatively, a moveable mirror can be shifted into the beam path guiding the light to a methanol-filled cuvette used as calibration sample. To achieve a sufficient working distance, a Zeiss Plan Neofluar 20x/0.5 NA objective is used, which results in a spatial resolution of 1 μm in the lateral plane and approximately 5 μm in the axial direction. For high-resolution images we apply a Zeiss water-immersion C-Apochromat 40x/1.2 NA objective providing subcellular resolution of 0.4 μm in the lateral plane and approximately 1 μm in the axial direction. Similar resolutions have been demonstrated to be possible for Brillouin microscopy in^65, 66^already. However, using high-NA objectives leads to a decrease of the Brillouin shift of 1 – 2 % for the objectives used here^67^. The sample is furthermore illuminated with a white light source to acquire a brightfield image after the Brillouin measurement. The Brillouin scattered light is collected in backscattering direction and separated from the brightfield illumination by a shortpass mirror (Thorlabs DMSP650) with a 650 nm cut-off wavelength. To achieve confocality, a second single-mode fiber guiding the light into the Brillouin spectrometer is used. In the spectrometer, the light is collimated and subsequently filtered by a band-pass (Thorlabs FBH780-10) to block Raman scattered light. The molecular absorption cell (Precision Glass Blowing TG-ABRB-I85-Q) absorbs light with the same frequency as the excitation laser (eg. Rayleigh scattered and reflected light)^68^. After passing the absorption cell, the beam is guided to two VIPA interferometers (Light machinery OP-6721-6743-4) with 15.2 GHz free spectral range. The interferometers lead to an angular dispersion of light with different frequencies^62^, which allows to image the Brillouin spectrum of the probed measurement point on the sCMOS camera (Andor Zyla 4.2 PLUS). For this purpose, we recorded two spectra per measurement point with an acquisition time of 500 ms each, which we found optimal in order to minimize the measurement uncertainty. The resulting random error amounted to 14MHz which is similar to values from other publications^51, 65^. The standard deviation of the mean value of measurements with independent calibrations was 10MHz (see also Supplementary Fig. 2). A spatially resolved map of the Brillouin shift was acquired by automatically translating the specimen with a motorized stage.

The obtained Brillouin images reveal the different viscoelastic properties of varying anatomical structures within living zebrafish larvae. We find that the Brillouin shift of the spinal cord increases during development. After spinal cord injury, we detect a significant, but transient drop of the Brillouin shift at the lesion site. Thus, our experiments demonstrate for the first time that zebrafish spinal cord tissue, mechanically mapped inside living and intact animals, presents distinct mechanical signals to mechanosensitive cells during development and regeneration. We also show explicitly that the influence of refractive index changes within the perispinal area of zebrafish larvae can be neglected when calculating the longitudinal moduli from the measured Brillouin signal, and that there are significant differences between tissues in *situ* and after dissection. This work constitutes the methodical basis to identify key determinants of mechanical central nervous system tissue properties and to test the relative importance of mechanics in combination with biochemical and genetic factors during biologically important processes *in vivo.*

## Results

### Brillouin microscopy mechanically characterizes anatomical structures in living zebrafish larvae

To identify anatomical structures and assign viscoelastic properties to distinct organs and tissues in the living zebrafish larva, we compared images acquired with Brillouin microscopy with images from brightfield and confocal fluorescence microscopy of the same specimens at 4 days post-fertilization (dpf) (Fig. 2). For this purpose, individual zebrafish larvae were anesthetized and immobilized using low gelling point agarose in a glass bottom dish suitable for optical imaging. Using Brillouin microscopy, we measured the mechanical properties of the larval tissues in all three anatomical planes. All specimens were subsequently imaged using brightfield microscopy, and fixed and stained with TRITC-conjugated phalloidin and DAPI for confocal fluorescence microscopy. While the formation of melanophores could be suppressed successfully, all zebrafish exhibited the two other types of pigment cells, xanthophores and iridophores. The latter can be observed as dark spots in all brightfield images (Fig. 2a,b) and caused reflections during the Brillouin image acquisition (Fig. 2d,e,f, Supplementary Note 1). The corresponding values were therefore discarded and displayed as white pixels in the Brillouin images.

**Figure 2.**
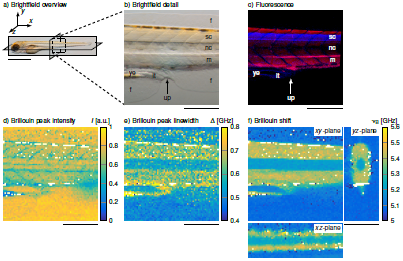
Images of the same zebrafish larva at four days post-fertilization (dpf). Brightfield and Brillouin images were obtained in a living specimen, fluorescence microscopy was performed after fixation following Brillouin microscopy, a) Brightfield image showing the specimen and the three anatomical planes. Scale bar, 1mm. b) Enlarged brightfield image of the region of interest containing the spinal cord (sc), the notochord (nc), muscle segments (m), the fins (f), the yolk extension (ye), the intestinal tissue (it) and the urogenital pore (up), c) Corresponding fluorescence image depicting the same structures – cell bodies in the spinal cord (nuclei, blue), muscle tissue (actin, red) and the notochord. Brillouin microscopy yields d) the peak intensity, e) the peak linewidth given as full width at half maximum and f) the Brillouin shift. The peak intensity detects anatomical structures that scatter or absorb light in the beam path. The peak width is indicative of the viscous properties. The Brillouin shift is displayed for the three anatomical planes indicated in a) and is proportional to the elastic properties of the sample. Scale bars b) – f), 200 μm.

In both the brightfield images (Fig. 2a,b) and the Brillouin peak intensity map (Fig. 2d), image contrast is governed by absorption and scattering processes in the specimen, which allows to discern prominent anatomical structures such as the notochord and the diagonal pattern of the muscle segments. The fluorescence image (Fig. 2c) allows additionally to distinguish the spinal cord tissue (blue) from the surrounding muscle tissue (red). While the aforementioned imaging modalities convey information about the presence of certain anatomical features, the Brillouin shift and the linewidth (full width at half maximum) show the mechanical properties of these features. The Brillouin peak linewidth depends on the phonon lifetime and is therefore directly related to the loss modulus and the viscosity of the sample (see Eq. [2]). For muscle and spinal cord tissue, the Brillouin peak linewidth (Fig. 2e) is higher than for the intestinal tissue and the notochord. The Brillouin shift depends on the longitudinal (storage) modulus and is indicative of the elastic, solid-like sample properties (see Eq. [1]). Our measurements (Fig. 2f) revealed a higher Brillouin shift of the yolk extension, the muscle and spinal cord tissue as compared to the intestinal tissue and notochord. The zebrafish notochord is formed by an outer layer of extra-cellular matrix (ECM) and sheath cells that surround an inner layer of vacuolated cells^69, 70^. Mechanically, the notochord has been described as a rigid, fluid-filled cylinder under pressure^70, 71^. Muscle tissue, nervous tissue and yolk, on the other hand, have all been described as viscoelastic materials exhibiting both fluid-like viscous properties and solid-like elastic properties^13, 72–76^. It is therefore not surprising that the solid-like characteristics displayed in the Brillouin shift maps are the lowest for the notochord as compared to the other types of tissue. The low Brillouin shift values for the intestinal tissue are presumably indicative of the presence of intestinal fluid that has been described as a Newtonian fluid with a negligible elastic contribution from the intestinal mucus^77^. The Brillouin shift map also revealed the diagonal pattern of the muscle segments and we were able to detect the fins in the *yz*-plane. After the measurements the larvae were released from the agarose and behaved normally after cessation of sedation.

### Brillouin shifts of the larval zebrafish spinal cord increase during development and decrease after injury

To detect changes of the mechanical properties of the spinal cord tissue during development, we probed the same specimens (*N* = 6) on three consecutive days. This allowed us to follow the evolution of the developmental mechanical phenotype that might serve as a signaling cue for neurons and glial cells residing in the tissue. Both the brightfield image (Fig. 3a) and the Brillouin peak intensity map (Fig. 3b) display the position of the spinal cord, notochord and the regular diagonal pattern of the muscle tissue. The position and contrast of these structures remained similar during the investigated time interval, whereas the Brillouin shift (Fig. 3c) of the spinal cord tissue increased from 3 dpf to 5 dpf.

**Figure 3.**
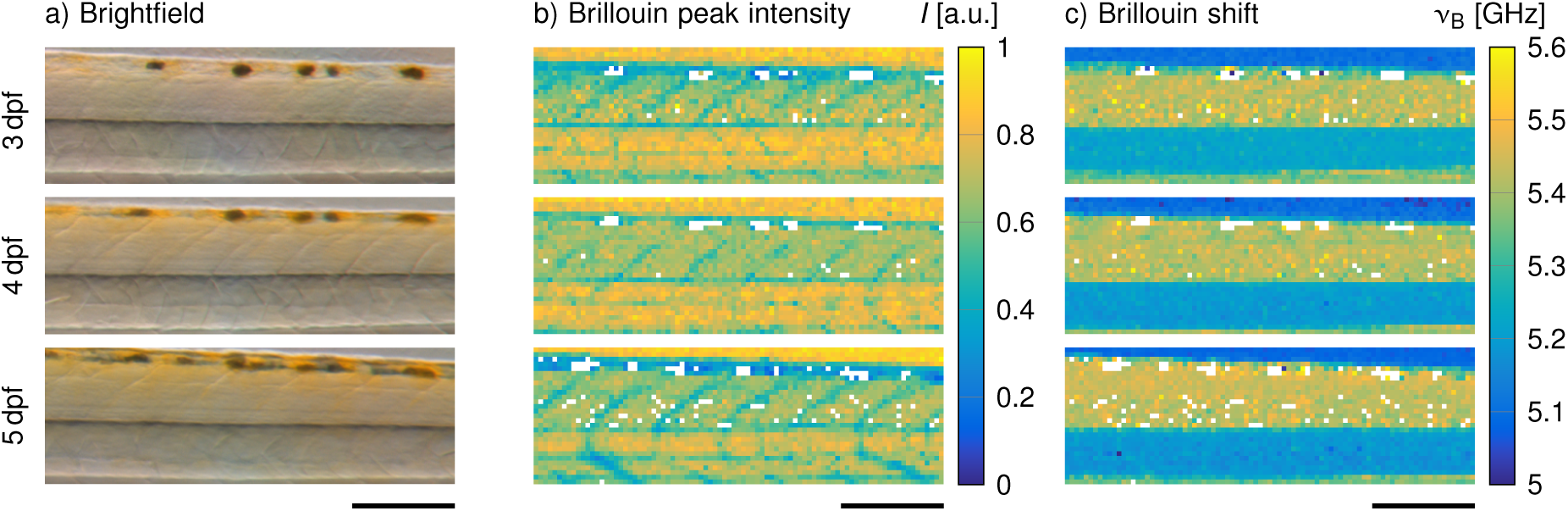
Brillouin measurements of a representative zebrafish larva during development, a) Brightfield images obtained directly after the Brillouin measurements, b) Corresponding peak intensities and c) Brillouin shifts of the same specimen. Scale bars, 150 μm.

To test the hypothesis that the regenerating zebrafish spinal cord changes its mechanical properties and thereby provides mechanical cues during regeneration, we acquired Brillouin images after spinal cord transection until 2 days post-injury (Fig. 4). All specimens (*N =* 6) were imaged immediately before the injury at 3 dpf, 1 h after the injury (3^+^ dpf) and on the two consecutive days (4 dpf and 5 dpf). The disrupted structure of the lesioned spinal cord was visible in both the brightfield image (Fig. 4a) and the Brillouin peak intensity map (Fig. 4b) where the lesion site displayed lower values as compared to neighboring spinal cord tissue. Similarly, the map of the Brillouin shift (Fig. 4c) showed lower values in the lesion site than in the surrounding tissue parts. It should be noted that the Brillouin shift of the lesioned area was still higher than that of the surrounding agarose gel. This indicated that there was not trivially just a liquid-filled gap but some sort of tissue with low elastic mechanical properties. As the repair of spinal cord progressed, the Brillouin shift increased gradually.

**Figure 4.**
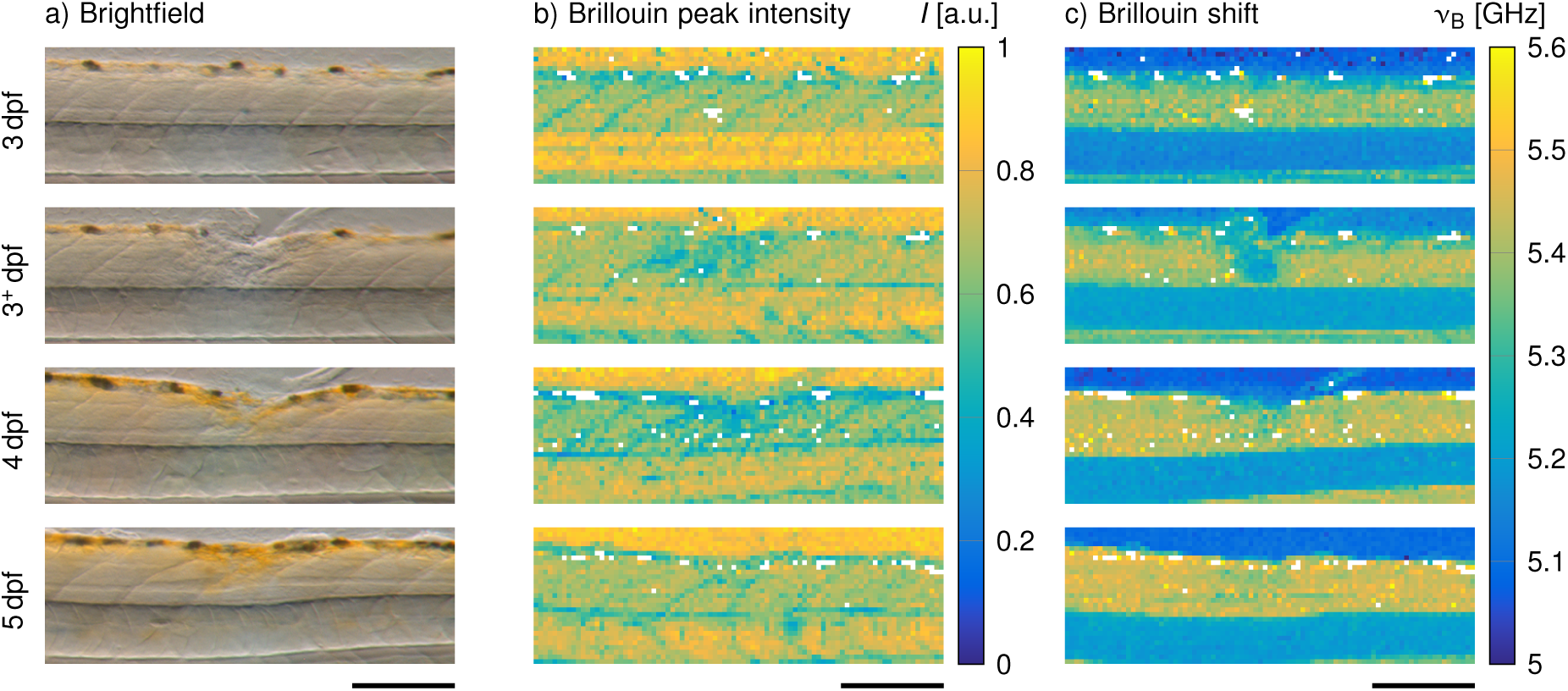
Brillouin measurements of a representative zebrafish larva after spinal cord injury. Lesioned larvae were probed at 3 dpf immediately before and 1 h after the injury (3+ dpf) as well as on the two subsequent days (4 dpf and 5 dpf). a) Brightfield images obtained directly after the Brillouin measurements, b) Corresponding peak intensities and c) Brillouin shifts of the same specimen. Scale bars, 150 μm.

The investigated time interval of lesioned animals coincided with the developmental period of uninjured animals and therefore allowed for direct comparison. To quantify the change of the Brillouin shift, we averaged the Brillouin shift values for three selected regions. The regions of interest in the uninjured animals correspond to the entire spinal cord tissue (Fig. 5a, red box). For the lesioned fish, the spinal cord tissue was divided into regions containing the lesion site (Fig. 5a, blue box) and regions for the remaining tissue (Fig. 5a, green box). The increase of the Brillouin shifts in uninjured animals from 3 dpf to 5 dpf (Fig. 5b) was significant (*p* = 0.0173). Brillouin shifts obtained from larvae after spinal cord injury (SCI) showed the same trend at regions next to the lesion site (Fig. 5c). The lesion site itself (Fig. 5d) displayed a significant decrease of Brillouin shifts immediately after SCI (*p* = 0.0021) which recovered to levels prior to the injury within the following two days. However, Brillouin shifts of the injured site at two days post-injury (5 dpf) did not reach levels next to the injury site, nor the levels of uninjured specimens of the same age (*p* = 0.0173).

**Figure 5.**
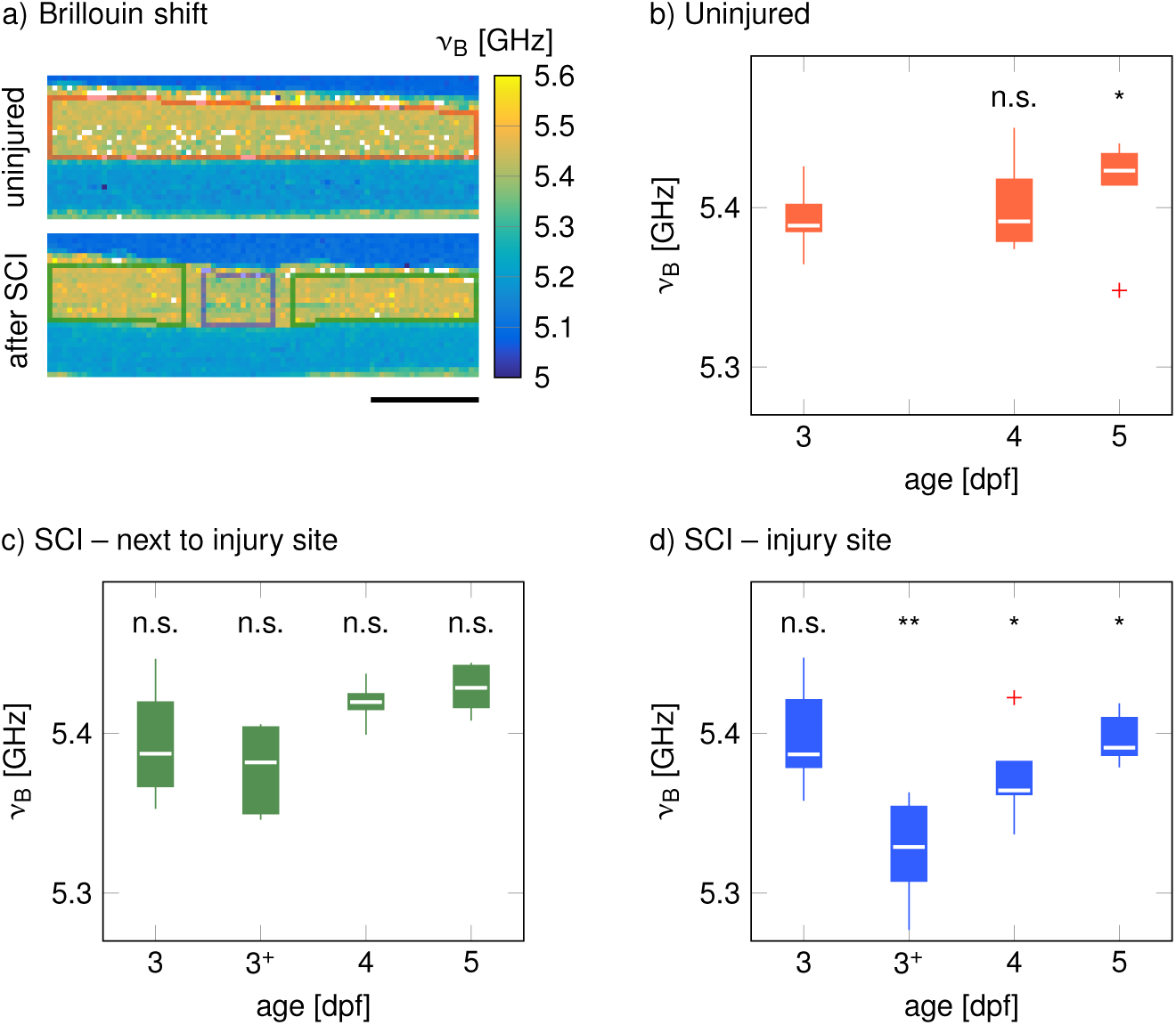
Quantification of the temporal development of the Brillouin shift during development and repair, a) The selected region of interest in the uninjured zebrafish (*N* = 6) comprises the complete spinal cord tissues (red box). For lesioned zebrafish (*N* = 6), the spinal cord tissue was divided into a region containing the lesion site (blue box) and a region for the remaining tissue (green box). Scale bar, 150 μm. b) – d) Box plots of the average Brillouin shifts of the respective areas in individual fish. The boxes indicate the interquartile ranges (IQR), the whiskers extend to the most extreme data still within 1.5 IQR of the respective quartile (Tukey boxplot) and the medians are depicted as white lines. Outliers are indicated as red crosses. Significance levels shown in b) refer to pairwise comparisons with 3 dpf. Indicated significant levels in c) and d) refer to pairwise comparisons of respective time points with uninjured animals in b). Brillouin shifts measured 1 h after spinal cord injury (3+ dpf) were also compared to 3 dpf values from uninjured specimens. \**p* < 0.05; \*\**p* < 0.01.

### A decrease of Brillouin shifts coincides with lower cell body densities

The mechanical properties of nervous tissues have been correlated to local cell body density and distribution using the presence of nuclei as a proxy for cell bodies^11, 46, 56^. To investigate if the changes of Brillouin shifts observed in the present study coincide with an altered tissue structure and cell body arrangement, we fixed and stained uninjured and lesioned zebrafish after Brillouin microscopy (Fig. 6). Uninjured spinal cord tissue displayed a packing of cell nuclei that was translationally invariant along the anterior-posterior axis of the spinal cord. This highly ordered arrangement corresponded to evenly distributed Brillouin shifts. As described above, the lesion site displayed significantly decreased Brillouin shifts in comparison to uninjured animals. The low values in the lesion site corresponded to a disorganized arrangement of cell nuclei and a lower fluorescence intensity of the DAPI signal as compared to neighboring, structurally intact tissue segments.

**Figure 6.**
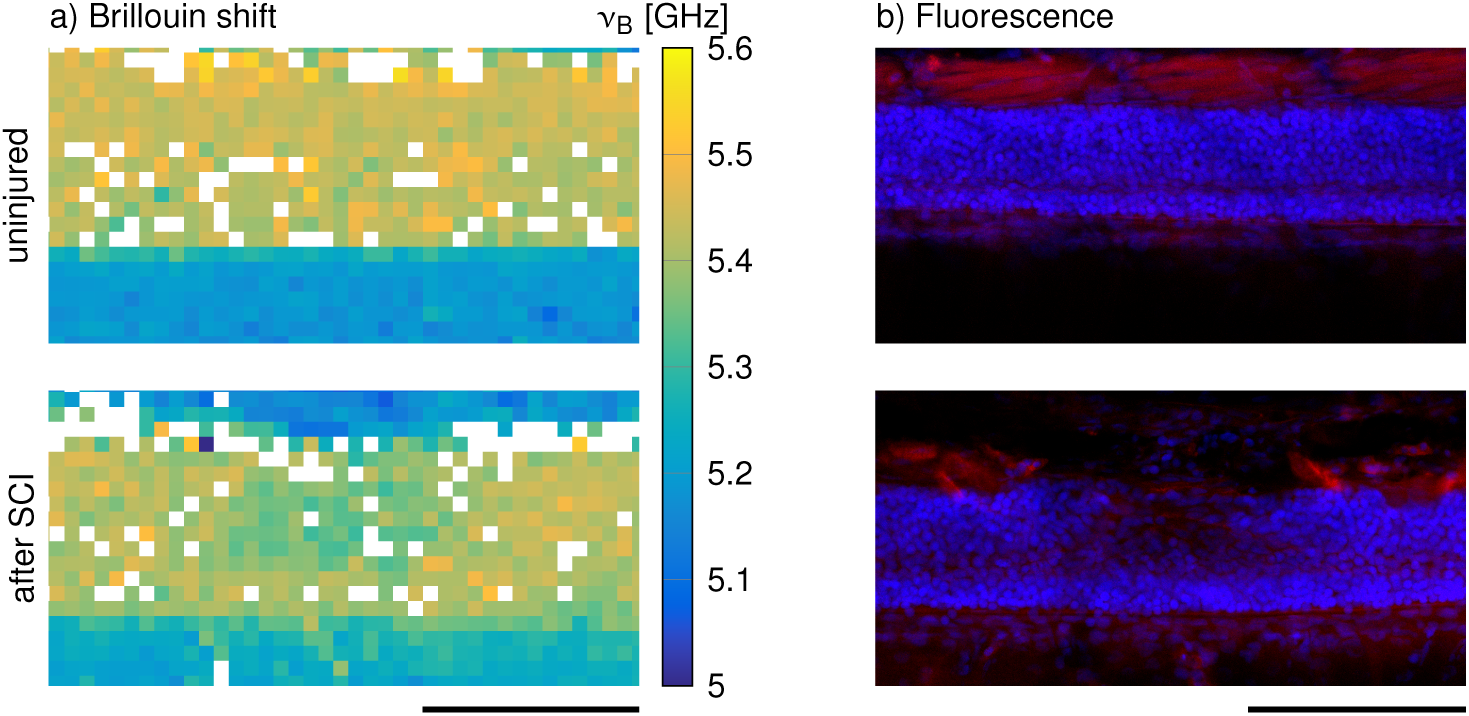
Comparison of a) the Brillouin shift and b) the corresponding fluorescence image for an uninjured and a lesioned zebrafish spinal cord at 5 days post-fertilization. The fluorescence image shows cell bodies (nuclei, blue) and muscle tissue (actin, red). SCI = spinal cord injury. Scale bars, 100 μm.

### Comparison of *in vivo* and *ex vivo* measurements

Previous publications addressing the mechanical characterization of muscle and nervous tissue reported greater elasticity values of muscle tissue in comparison to spinal cord tissue^76, 78, 79^. Our Brillouin measurements, however, did not reveal a discernible mechanical difference between muscle and spinal cord tissue. To investigate this apparent discrepancy, we obtained high-resolution Brillouin images in the *yz*-plane of the perispinal area in anesthetized zebrafish larvae. These measurements, too, show a similar Brillouin shift of spinal cord and muscle tissue in living zebrafish larvae (Fig. 7a). Since the aforementioned studies employed contact-based methods to probe mechanical tissue properties and therefore required tissue dissection and possibly tissue-sectioning, we suspected that the reason for the reported presence of mechanical differences between muscle and spinal cord tissue might originate from the differing sample preparation. To investigate the influence of sample preparation on the mechanical tissue properties in larval zebrafish, we obtained Brillouin images in the xy-plane of the perispinal area in transverse zebrafish tissue slices. These images showed greater Brillouin shifts in the muscle tissue as compared to the spinal cord tissue (Fig. 7b). Within the spinal cord tissue, we detected that areas that predominantly comprise cell bodies (Fig. 7c, blue) displayed higher Brillouin shifts than areas that contain mostly axons (Fig. 7c, green). The observed difference of mechanical properties between spinal cord areas and muscle tissue *ex vivo* could also be confirmed with AFM indentation measurements (see Supplementary Note 2 and Supplementary Fig. 3) and might be attributable to multiple factors. First, the acute zebrafish sections were kept in a medium of which the chemical composition was tailored to the nutritional demand of acute zebrafish spinal cord tissue^80, 81^, but not muscle tissue. This might lead to the faster onset of decomposition in muscle tissue than spinal cord tissue and thus, might serve to explain the relative changes of mechanical tissue properties. Second, muscle tissue *post-mortem* also exhibits a chemically induced stiffening, *rigor mortis*. To test if we detect time-dependent mechanical changes of muscle tissue *post-mortem* that might reflect the onset of *rigor mortis*, we performed Brillouin measurements of the perispinal area in sacrificed, but not sectioned, zebrafish larvae. The data showed that muscle tissue increasingly stiffened with time *post-mortem*, while spinal cord tissue, under the here described conditions, exhibits only small changes of mechanical tissue properties during the investigated time interval (Supplementary Note 3 and Supplementary Fig. 4). Third, vibratome-sectioning releases tension in both muscle and spinal cord tissue, which might contribute to the differences observed *ex vivo*. Also, the different observation directions between *in vivo* and *ex vivo* measurements could also influence the measured Brillouin signals.

**Figure 7.**
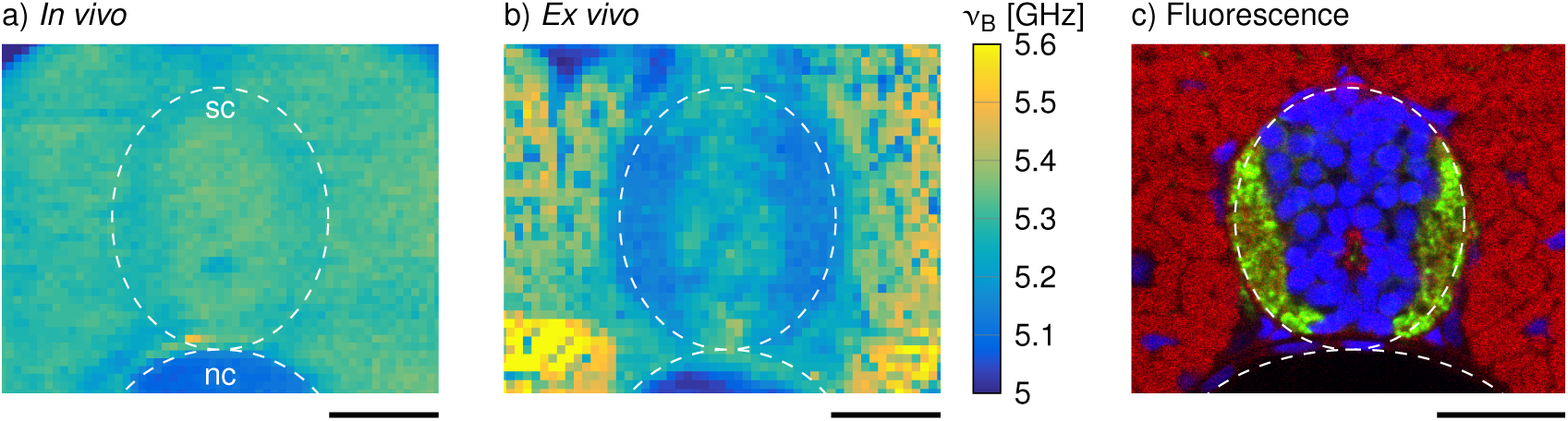
Representative high-resolution Brillouin images of the perispinal area in a) an anaesthesized zebrafish larva and b) an acute tissue slice from a vibratome-sectioned specimen. c) Immunohistochemistry of a transverse zebrafish section showing fluorescent labeling of cell bodies (nuclei, blue) and axons (acetylated tubulin, green) in the spinal cord as well as muscle tissue (actin, red) in the surrounding area. The approximate positions of the spinal cords (sc) and the notochords (nc) are indicated by dashed lines. Scale bars, 25 μm.

And finally, neglecting the contribution of the refractive index could also conceal differences of the mechanical properties when calculating the longitudinal modulus from the measured Brillouin shift.

### Relation between Brillouin signals and mechanical properties for larval zebrafish tissues

In order to calculate the actual longitudinal modulus and the viscosity of a transverse zebrafish section from the measured Brillouin shifts and linewidths (see equations [1] and [2] in Methods), we acquired the refractive index distribution (Fig. 8a) using quantitative phase imaging and calculated the density (see Supplementary Note 4) for the perispinal tissues. We confirmed the basic validity of this approach by analyzing ideal elastic (hydrogel) and viscous (glycerol-water mixtures) test samples with Brillouin microscopy and AFM-based nanoindentation and a falling-ball viscosimeter, respectively (see Supplementary Note 5). The maps of Brillouin shift *V*_*b*_ and linewidth Δ and of the resulting longitudinal modulus *M′* and viscosity *η* are shown in Fig. 8 b,c and Fig. 8 d,e, respectively. Our data show that the longitudinal modulus of the spinal cord is lower as compared to muscle tissue. Within the spinal cord, regions containing axons show a lower modulus than regions that contain mostly cell bodies. In fact, the longitudinal modulus map (Fig. 8d) reflects the information visible in the Brillouin shift map (Fig. 8b), indicating very little, if not negligible, contribution of the refractive index. The same holds true for the relation between the linewidth (Fig. 8c) and the viscosity (Fig. 8 e).

**Figure 8.**
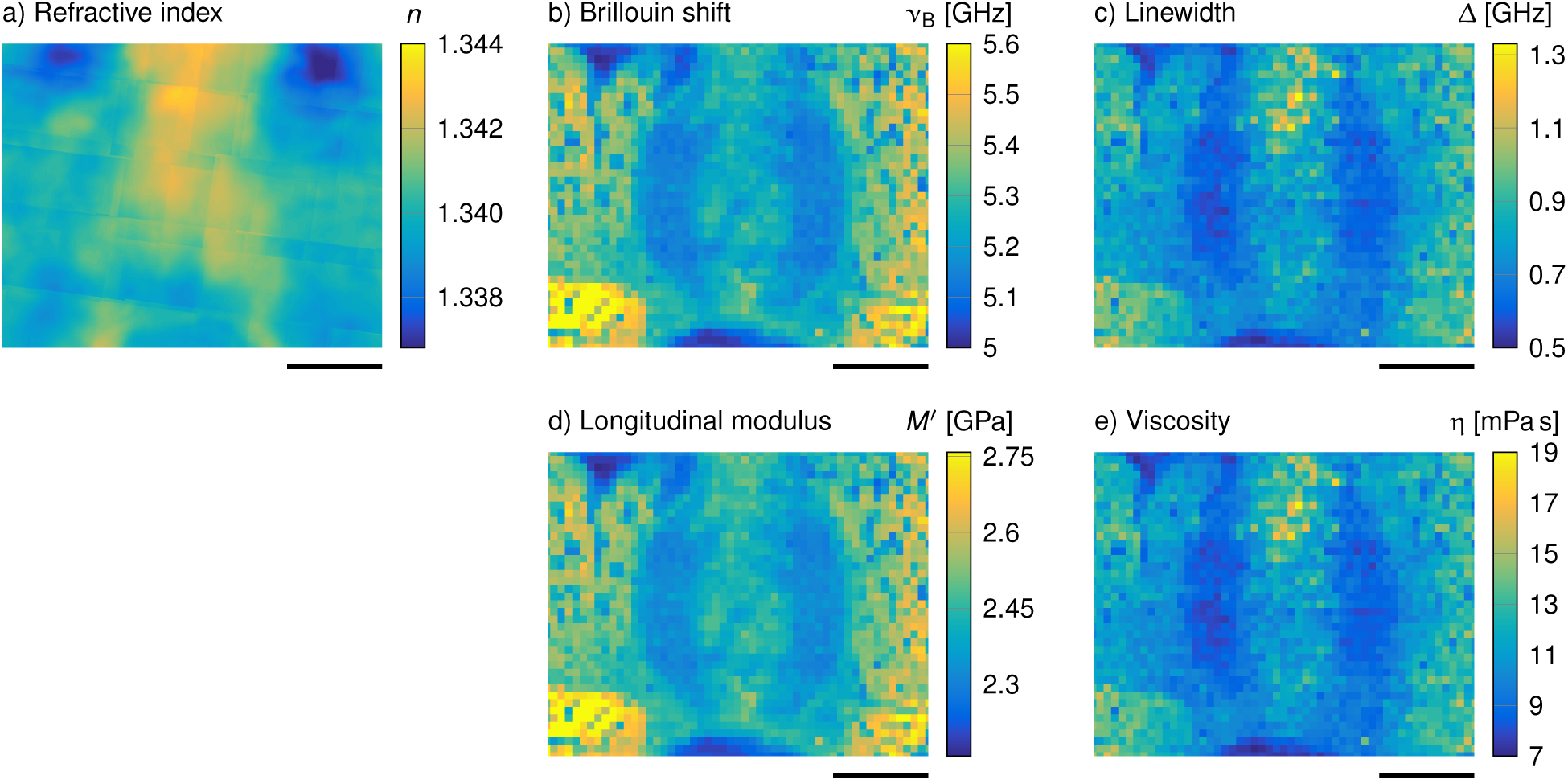
Determination of the longitudinal modulus *M′* and the viscosity *η* from the refractive index *n*, the Brillouin shift Vb and the linewidth Δ for larval zebrafish tissue, a) Map of the refractive index *n* obtained by quantitative phase imaging, b) Brillouin shift and c) linewidth obtained with the Brillouin microscope for an acute zebrafish slice, d) Longitudinal modulus *M′* and e) viscosity *η* determined with Eq. [1] and [2] from b) and c). Scale bars, 25 μm.

On the assumption that the refractive index is comparable in living zebrafish larvae and acute sections, we can conclude that the Brillouin shift as well as the Brillouin peak linewidth correlate directly with the longitudinal modulus and the viscosity, respectively. Hence, the Brillouin signals reported in this study constitute mechanical properties. Consequently, the changes of the Brillouin shift of the spinal cord tissue during development and regeneration after injury correspond to changes of the mechanical tissue properties that potentially present mechanical signals to mechanosensitive cells residing in the respective tissues.

## Discussion

In this report, we demonstrated the first systematic investigation of the mechanical properties of larval tissues in living zebrafish. By using a custom-built confocal Brillouin microscopy setup, we were able to measure the mechanical phenotypes of distinct anatomical structures. The measurements show a lower Brillouin shift and a lower Brillouin peak linewidth of the notochord and intestinal tissue as compared to the spinal cord, muscle tissue and yolk extension. This conforms with previous studies describing the notochord and the intestinal tissue as fluid-filled structures^69^, and spinal cord, muscle tissue and yolk as viscoelastic materials^13, 72–76^. As for the yolk extension, our measurements yielded comparable values as previously reported for the yolk in zebrafish embryos^60^. Our results furthermore showed that developmental and pathological processes of larval zebrafish spinal cords coincide with significant Brillouin shift changes. During development, we detected an increase of the Brillouin shifts. After injury, the Brillouin shifts measured in the lesion site decreased immediately, but gradually increased thereafter.

The data also show that the Brillouin shifts of muscle and spinal cord tissue exhibit comparable values *in vivo*. This seemingly contrasts previous results using contact-based techniques like AFM-based indentation measurements^76, 78, 79^. However, AFM measurements are performed using *ex vivo* samples. For vibratome-sectioned zebrafish tissues, Brillouin measurements show different Brillouin shift for the different types of tissue. In fact, the Brillouin shift of the muscle tissue is higher than the shift of the spinal cord tissue, which compares to the relative mechanical properties assessed by indentation measurements. The parameters that contribute to mechanical tissue properties *in vivo* as well as *ex vivo* could be manifold. For instance, indentation measurements require exposed sample surfaces and possibly sample sectioning. This leads to necrosis, apoptosis, the release of tension and hydrostatic pressure changes which all may induce alterations of the mechanical tissue properties. On the other hand, the acquisition of high-resolution images of living organisms using confocal microscopy requires sample immobilization and complete lack of muscle twitching. In zebrafish, this sedation is commonly achieved by immersion in tricaine (MS222) containing media^82^. Tricaine might induce acidification in weakly buffered solutions^82^. Acidification has been shown to influence the stiffness of rat brain tissue^83^ and might similarly act on the mechanical properties of zebrafish nervous tissue. Furthermore, the larval tissue sections were incubated in a solution whose chemical composition was tailored to the nutritional demand of zebrafish nervous tissue, but not muscle tissue. This might induce a faster decomposition of the muscle tissue and/or necrosis. The difference between mechanical differences found *ex vivo*, that were absent *in vivo*, emphasizes the need for methods, such as Brillouin microscopy, to quantitatively map mechanical properties inside living animals.

The Brillouin shift scales with the longitudinal modulus, the refractive index and the density of the sample (see Eq. [1]). For sectioned tissues, we determined the refractive index distribution and calculated the longitudinal modulus and viscosity. In fact, the longitudinal modulus map reflects the information visible in the Brillouin shift map, indicating very little, if not negligible, contribution of the refractive index. Assuming that the amplitude of refractive index variations within the sample is similar between *in vivo* and *ex vivo* measurements, the Brillouin shift directly corresponds to the longitudinal modulus. However, this assumption might not hold true for other samples. Recent advances in refractive index tomography enable to measure the three-dimensional refractive index distributions for thin samples and single cells^84, 85^. Hence, by combining refractive index tomography and Brillouin microscopy, an unambiguous distribution of the density and the elastic and viscous properties could be obtained for these samples as well.

Still, the interpretation of the acquired longitudinal modulus remains challenging. Brillouin microscopy acquires information on very short time and length scales, whereas AFM provides information for larger time and length scales. Hence, longitudinal and Young’s modulus must not necessarily correlate, but could complement each other by providing information about different temporal and spatial regimes. Furthermore, it has been proposed that Brillouin microscopy is largely sensitive to the water content of the probed volume^86^. While we also found a correlation between Brillouin shift and density of cell nuclei, what other constituents of living biological tissue Brillouin scattering is really sensitive to remains an interesting area of exploration in the future. In any case, the changes of the mechanical properties observed in the larval zebrafish may serve as signaling cues for mechanosensitive neural cell types residing in the spinal cord tissue. In fact, the functional recovery of zebrafish after injury, and the lack thereof in mammals, is associated with cell types and cellular processes that are mechanosensitive both *in vitro* and *in vivo*.

This work constitutes the methodical basis to identify key determinants of mechanical central nervous system tissue properties and to test the relative importance of mechanics in combination with biochemical and genetic factors during biologically important processes *in vivo*. Future insights into the mechanical mechanisms promoting or inhibiting neuronal regrowth, enabled by Brillouin microscopy as demonstrated in this study, might therefore constitute important additions to biochemical, or cell and molecular biological analyses, and could eventually even provide novel therapeutic strategies for affected humans.

## Methods

### Animals

All zebrafish were kept and bred in the fish facility of BIOTEC, TU Dresden as described in^87^. Embryos from wild-type (WIK) zebrafish were incubated in PTU (1-phenyl-2-thiourea, Sigma-Aldrich, 0.00375% in E3 embryo medium) after 1 dpf to prevent melanogenesis^88^.

### Mounting of zebrafish larvae for in vivo Brillouin and brightfield imaging

Larvae were anesthetized in MS222 (0.01% in E3, Sigma-Aldrich) and placed in a lateral position on a 30 μm thick, flat polyacrylamide (PAA) gel that had been polymerized on a glass bottom dish suitable for optical imaging. A drop ( 200 μl) of low gelling point agarose (1% in E3, 30°C, Sigma-Aldrich) was used to immobilize the larvae in this position for subsequent imaging. Immobilized larvae were then immersed in MS222 (0.01% in E3) during imaging. All specimens were released from the agarose embedding between Brillouin measurements and kept under standard conditions^87^.

### Fluorescence microscopy

After obtaining the final Brillouin image, zebrafish larvae were sacrificed by an overdose of MS222, fixed with PFA (4% in phosphate buffered saline (PBS)), subjected to multiple washing and permeabilization steps using PBSTx (0.8% TritonX-100, 1% DMSO in PBS) and stained with DAPI and phalloidin-TRITC (both 1:1000 in PBS). Larvae were then washed with PBS, incubated in glycerol (75% in PBS) for 3 h and mounted in DABCO containing mounting medium (2.5% DABCO, Sigma-Aldrich, 90% glycerol in 50mM Tris-HCl, pH 8).

For immunohistochemistry on vibratome-sectioned zebrafish larvae, 4 day old specimens were sacrificed and chemically fixed as described above. The larvae were then embedded in 2.5% low gelling point agarose and sectioned at 150 μm to 200 μm. Thereafter, sections were subjected to multiple washing and permeabilization steps using PBTx for approximately 2 h before incubation in blocking buffer (4% BSA in PBSTx) for 1 h. Sections were then incubated with a primary antibody against acetylated tubulin (Sigma T6793, 1:500 in blocking buffer) overnight at 4 °C. Afterwards, sections were washed twice in PBSTx for consecutive 5 min, 10 min, 15 min and 30 min before incubation with phalloidin (conjugated to Alexa Fluor 647, Invitrogen A22287, 1:1000) and secondary antibody (goat anti-mouse Alexa Fluor 488, Invitrogen A11001, 1:200) in blocking buffer overnight at 4 °C. The next day, sections were washed twice in PBSTx for consecutive 5 min, 10min, 15 min and 30min before washing twice with PBS, counterstaining with DAPI and mounting in DABCO-containing mounting medium.

### Image acquisition and processing

Brightfield images were obtained using the Olympus Stereomicroscope SZX 16 and the SDF PLAPO 0.8x/0.12 NA and 2.0x/0.3 NA objectives. Confocal fluorescence microscopy was performed to image fixed and stained larvae using the Zeiss LSM700 and a Plan Apochromat 10x/0.45 NA objective. The fluorescence images displayed in this study are maximum z-projections of four optical planes (≈ 1μm each) which correlates to the section thickness measured by Brillouin microscopy. Brightfield and fluorescence images were adjusted in brightness and contrast using Fiji and assembled with CorelDraw.

### Spinal cord lesion

Larvae were anesthetized in MS222 (0.01% in E3) and immobilized as described for Brillouin imaging. Lesions were performed manually at 3 dpf by inserting the broken tip of a pulled injection glass capillary in the most ventral part of the spinal cord at the level of the urogenital pore, and moving the tip dorsally in a quick motion. Notochords and major blood vessels remained intact. Larvae were subsequently released from the agarose embedding.

### Brillouin scattering

Brillouin scattering is an inelastic scattering process between the incident light and periodic mass density fluctuations due to travelling sound waves with sound velocity *V* within the sample. This results in an upward and downward frequency shift *V*_*b*_ (i.e. Brillouin shift) of the incident light, which is given by^50, 89^

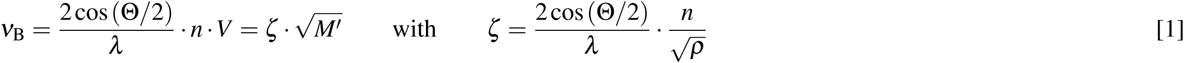

where Θ is the scattering angle and *λ* the wavelength. The mechanical properties of the sample are described by the longitudinal modulus *M*′, the refractive index n and the mass density *ρ* of which the refractive index and the mass density are directly coupled for most biological samples^84, 90–93^. The linewidth Δ (full-width at half maximum) of the Brillouin peak is given by_94, 95_

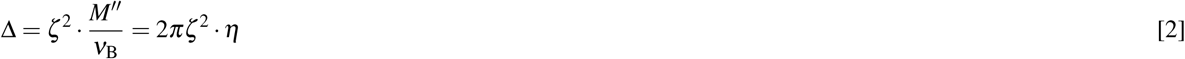

where *M′′* is the loss modulus and *η* the viscosity.

### Brillouin imaging

The acquisition of the camera images as well as the control of the translation stage and the objective lens’ z-position are performed using a custom MATLAB program. The acquired raw data are stored as a HDF5 file allowing for efficient data handling^96^. The subsequent data analysis also uses a custom MATLAB program and is done in multiple steps. First, the one-dimensional Brillouin spectra are extracted from the two-dimensional camera images. The position, width and intensity of the Brillouin and Rayleigh peaks are then determined by fitting Lorentzian functions to every peak. In order to calibrate the frequency axis, a reference measurement of a methanol sample placed in the calibration beam path is acquired every 10 min by moving a mirror into the beam path (Fig. 1). This allows to obtain an absolute value for the Brillouin shift and compensates for possible drifts of the VIPA setup during the measurement. For every reference measurement we fit the theoretical, non-linear frequency axis to the Brillouin spectrum according to^97^. Interpolating along the temporal axis yields a calibrated frequency axis for every time point during the measurement, which is used to calculate the Brillouin shift in GHz. Plotting the calibrated Brillouin shift over the position of the translation stage finally leads to a spatially resolved map of the mechanical properties of the sample. All white pixels displayed in the maps correspond to coordinates in which the sample caused strong back-reflections due to large refractive index gradients or the Brillouin signal was absorbed due to strong pigmentation (See Supplementary Note 1). These values were discarded.

### Preparation of zebrafish sections

Zebrafish larvae were sacrificed by an overdose of MS222 (0.2% in PBS, Sigma-Aldrich), embedded in low gelling point agarose (2.5% in artificial cerebrospinal fluid, Sigma-Aldrich) and cut into transverse sections using a vibratome. The artificial cerebrospinal fluid (aCSF) contained (in mM) 134 NaCl, 2.9 KCl, 1.2 MgCl_2_, 2.1 CaCl_2_, 10 HEPES buffer, and 10 glucose, adjusted to pH 7.8 with NaOH^81^. A section thickness of 300 μm was used for indentation measurements as well as Brillouin microscopy. Refractive index (RI) measurements were performed using 200 μm thick sections. Zebrafish sections selected for indentation measurements were immobilized on tissue culture plastic (TCP) with Histoacryl (B. Braun) which was sparsely applied between the TCP and the agarose embedding at a distance from the zebrafish section. The sections were submerged in cooled aCSF during indentation measurements. For both Brillouin microscopy and RI measurements, the sections were immobilized by sandwiching them between two glass slides suitable for optical imaging, and surrounded by aCSF. All vibratome-sections were obtained from 4 day old larvae.

### Quantitative phase imaging

Quantitative phase and intensity images were recorded with a commercial phase-imaging camera (SID4-Bio, Phasics) attached to an inverted microscope (IX71, Olympus) as described in^98^. The imaged zebrafish section was larger than the field of view (FOV) covered by the camera sensor and thus we applied an image-stitching approach. Intensity images were recorded manually with a 40% to 80% overlap and background corrected by division with a reference image. Stitching was performed with an appropriate type of descriptors^99^ in the intensity images, applying a translational model, and isolating valid descriptors with a random sample consensus approach. The images were blended using the median value of the individual images. Quantitative phase images exhibit random phase offsets which were corrected with an iterative approach. In each iteration, the stitched phase image was assembled using the image locations obtained from stitching the intensity data. Then, the mean phase of each individual image, excluding a ten pixel border around the image, was corrected to match that of the corresponding FOV in the stitched image for the subsequent iteration. A total of 20 iterations were performed. The phase measured corresponds to the difference in the optical path difference through the zebrafish section. By taking into account the thickness of the section, the integral refractive index *n*(*x*, *y*) at each point of the section can be computed using

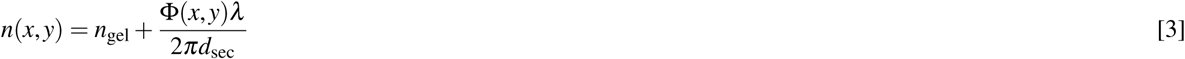

where *n*_gel_ = 1.3395 is the refractive index of the hydrogel measured with an Abbe refractometer (2WAJ, Arcarda), Φ(*x*,*y*) is the measured quantitative phase, *λ* = 647nm is the wavelength used for imaging and *d*_*sec*_ = 200 μm is the thickness of the zebrafish section.

### Statistical analysis

To compare the Brillouin shifts of different regions within the specimens’ spinal cords, the values corresponding to every region had to be identified. For this purpose, all measurement points belonging to the specimen’s spinal cord were selected first. For the injury site, all values in the spinal cord in an area with a width of 100 μm around the center of the lesion were chosen. The region next to the injury site included all values in the spinal cord separated from the injury site by a 25 μm clearance to exclude the transient values between both regions. All Brillouin shifts corresponding to the identified regions were averaged for the individual specimen, yielding *N* = 6 mean values for every region and for the injured and uninjured zebrafish, respectively. The mean values were subjected to statistical analysis by the two-tailed Mann-Whitney U-test using MATLAB and printed as boxplots. The shown asterisks indicate the significance levels: \**p* < 0.05 and \*\**p* < 0.01.

## Acknowledgements

We thank Prof. Matthias Kirsch from the Department of Neurosurgery at the University Hospital Carl Gustav Carus, Daniel Wehner from the Center for Regenerative Therapies Dresden, and James Briscoe from the Francis Crick Institute, London, for helpful discussions, the Advanced Light Microscopy Facility of the Max Delbriick Center for Molecular Medicine in the Helmholtz Association for technical support, the Center for Molecular and Cellular Bioengineering (CMCB) Fish Facility for supplying the zebrafish specimens and the CMCB Light Microscopy Facility (partly funded by the State of Saxony and the European Fund for Regional Development - EFRE) for technical support and provision of the confocal fluorescence microscope. Financial support from the Alexander-von-Humboldt Stiftung (Humboldt-Professorship to J.G.) and the European Commission through an ERC Starting Grant (”LightTouch”, grant agreement number 282060 to J.G.) and the BIOPOLITN (Marie Sklodowska Curie Action of the Horizon2020 program, grant agreement number 607350) is gratefully acknowledged.

## Author contributions statement

R.S. realized the Brillouin microscope and evaluated the Brillouin measurements. S.M. prepared the zebrafish specimens, acquired and evaluated the brightfield and fluorescence images, and performed and evaluated the AFM measurements on zebrafish tissues. R.S. and S.M. conducted the Brillouin measurements and wrote the manuscript with contributions from all authors. S.A. helped with sample preparation, prepared the polyacrylamide hydrogels, and performed and evaluated the corresponding AFM measurements. G.C., P.M. and K.K. conducted the refractive index measurements. C.M. and C.Z. supported the evaluation of the Brillouin measurements. R.S., S.M., J.C. and J.G. designed the experiments. All authors reviewed the manuscript.

## Additional information

Accession codes The datasets generated during and/or analyzed during the current study are available from the corresponding author on reasonable request. Competing financial interests The authors declare no competing financial interests.

